# SbcB facilitates natural transformation in *Vibrio cholerae* in an exonuclease-independent manner

**DOI:** 10.1101/2024.09.25.615017

**Authors:** Triana N. Dalia, Ankur B. Dalia

## Abstract

Natural transformation (NT) is a conserved mechanism of horizontal gene transfer in bacterial species. During this process, DNA is taken up into the cytoplasm where it can be integrated into the host genome by homologous recombination. We have previously shown that some cytoplasmic exonucleases inhibit NT by degrading ingested DNA prior to its successful recombination. However, one exonuclease, SbcB, counterintuitively promotes NT in *Vibrio cholerae*. Here, through a systematic analysis of the distinct steps of NT, we show that SbcB acts downstream of DNA uptake into the cytoplasm, but upstream of recombinational branch migration. Through mutational analysis, we show that SbcB promotes NT in a manner that does not rely on its exonuclease activity. Finally, we provide genetic evidence that SbcB directly interacts with the primary bacterial recombinase, RecA. Together, these data advance our molecular understanding of horizontal gene transfer in *V. cholerae*, and reveal that SbcB promotes homologous recombination during NT in a manner that does not rely on its canonical exonuclease activity.

**IMPORTANCE:** Horizontal gene transfer by natural transformation contributes to the spread of antibiotic resistance and virulence factors in bacterial species. Here, we study how one protein, SbcB, helps facilitate this process in the facultative bacterial pathogen *Vibrio cholerae*. SbcB is a well-known for its exonuclease activity (*i*.*e*., the ability to degrade the ends of linear DNA). Through this study we uncover that while SbcB is important for natural transformation, it does not facilitate this process using its exonuclease activity. Thus, this work helps further our understanding of the molecular events required for this conserved evolutionary process, and uncovers a function for SbcB beyond its canonical exonuclease activity.

## INTRODUCTION

Cells can acquire novel traits through horizontal gene transfer by natural transformation. During this process, cells take up free double-stranded DNA (dsDNA) from the environment. In Gram-negative bacteria, dsDNA is first taken up across the outer membrane via the combined action of dynamic DNA-binding type IV pili (1, 2) and the periplasmic DNA-binding molecular ratchet ComEA (3, 4). Then, a single strand of this DNA is translocated across the inner membrane by ComEC and ComF(C). Once this single-stranded DNA (ssDNA) is translocated into the cytoplasm, it can interact with proteins that help facilitate its integration into the host genome. This includes DprA, which protects ingested DNA and helps recruit RecA to initiate homologous recombination (5).

Because the tDNA that is imported into the cytoplasm is ssDNA, it is potentially sensitive to the action of ssDNA exonucleases that may act to degrade this DNA before it can be integrated into the genome. Indeed, we have previously found that deletion of the ssDNA exonucleases *recJ* and *exoVII* enhances NT in *V. cholerae* (6). In particular, Δ*recJ* Δ*exoVII* mutants can be transformed at high rates with tDNA containing short arms of homology (6), which is consistent with these proteins normally degrading the ends of ssDNA tDNA in the cytoplasm to inhibit NT.

Contrary to this negative effect of the RecJ and ExoVII ssDNA exonucleases on NT, our prior results suggest that one ssDNA exonuclease, SbcB, is required for optimal NT in *V. cholerae* (6). SbcB is one of the most well-studied bacterial exonucleases, primarily having been characterized in the model microbe *Escherichia coli*. It was originally identified biochemically and called Exonuclease I or ExoI (7), and subsequent genetic analysis uncovered the gene responsible and referred to it as SbcB (8) and XonA (9). In this study, we sought to determine how SbcB counterintuitively promotes NT in *V. cholerae*.

## RESULTS

### SbcB is required for optimal NT

The genes required for NT in *V. cholerae* are tightly regulated and induced via two distinct regulatory inputs: chitin oligosaccharides and quorum sensing (10, 11). Our prior experiments assessed the impact of *sbcB* on NT in a strain background that is dependent on both of these regulatory inputs. Thus, one possibility is that SbcB is important for regulating the expression of genes required for NT. Regulatory control of NT can be bypassed by ectopically expressing the master regulator of competence TfoX (10) and by constitutively activating quorum sensing via deletion of LuxO (1, 11, 12). So to test this hypothesis, we assessed the impact of *sbcB* on NT in a background where *tfoX* is ectopically expressed (*P*_*tac*_-*tfoX*) and *luxO* is deleted. In this genetic background, we found that Δ*sbcB* exhibited a ∼1.5-log decrease in NT (**Fig. 1**), which phenocopies what was previously observed when cells relied on natural induction via chitin and quorum sensing (6). This indicates that SbcB acts downstream of the regulatory steps that lead the expression of genes required for NT. All strains used throughout the remainder of the study contained P_tac_-*tfoX* and Δ*luxO* mutations, which allowed us to bypass the need for chitin induction to study the effect of SbcB on NT.

**Fig. 1.**
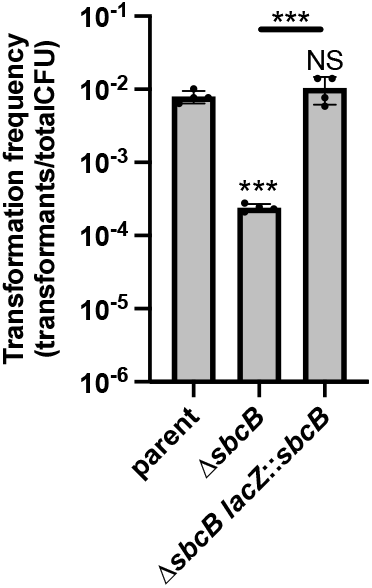
SbcB facilitates NT in *Vibrio cholerae*. Natural transformation assays of the indicated strains. Data are from four independent biological replicates and shown as the mean ± SD. Statistical comparisons were made by one-way ANOVA with Tukey’s multiple comparison test of the log-transformed data. NS, not significant. *** = *p* < 0.001

### SbcB does not facilitate the uptake of tDNA during NT

Next, we sought to identify the step of NT that is affected by SbcB. The first step of NT is the uptake of DNA from the environment, which requires the combined activities of dynamic DNA-binding competence type IV pili (T4P) to initiate the translocation of DNA across the outer membrane and a DNA-binding periplasmic molecular ratchet called ComEA. ComEA is a periplasmic dsDNA-binding protein, and it dynamically relocalizes upon DNA uptake, which can be monitored via functional fluorescent fusions like ComEA-mCherry (3, 4). The localization of ComEA-mCherry shifts from diffuse within the periplasm in the absence of tDNA uptake, to the formation of fluorescent foci when dsDNA is taken up into the periplasm (**Fig. 2A**). Thus, tracking the formation of ComEA-mCherry foci can allow for quantification of DNA uptake across the outer membrane. We assessed the formation of ComEA-mCherry foci after cells were incubated with tDNA for 30 mins (**Fig. 2A**). In the parent strain, almost all cells exhibited 1 or more ComEA-mCherry focus after this 30 min incubation (**Fig. 2B**). Consistent with tDNA uptake requiring T4P, mutating the critical T4P component PilQ (13, 14) prevents the formation of ComEA-mCherry foci (**Fig. 2B**). When we tested the *sbcB* mutant, we found that this strain could take up DNA into the periplasm in a manner that was indistinguishable from the parent (**Fig. 2B**; compare 30 min timepoints).

**Fig. 2.**
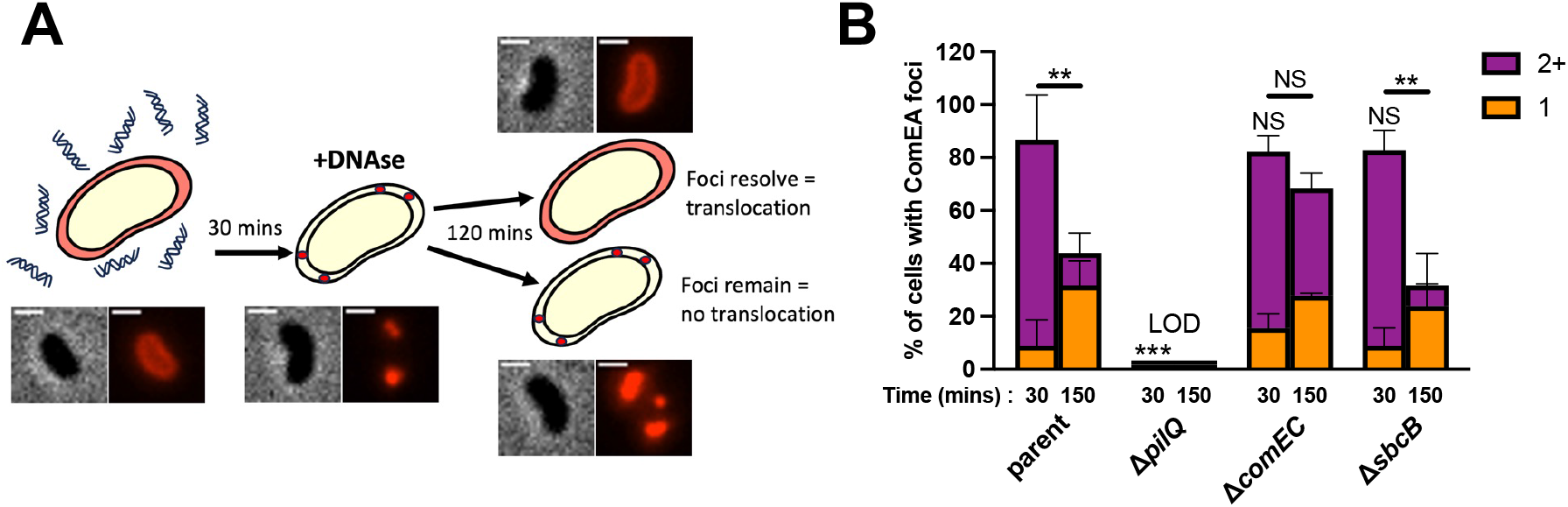
SbcB does not affect DNA uptake during NT. (**A**) Schematic and representative images of the experimental setup used to assess DNA uptake during NT. Cells contained a ComEA-mCherry fusion and were incubated with tDNA. The presence of a ComEA-mCherry focus indicates that tDNA is taken up into the periplasm. After 30 mins, DNAse was added to remove all extracellular DNA. The subsequent tracking of ComEA-mCherry foci was used as an indirect readout to demarcate the translocation of periplasmic tDNA into the cytoplasm. Scale bar, 1 µm. (**B**) The assay described in **A** was performed in the indicated strain backgrounds and the frequency of cells with 1 or more than 1 (2+) ComEA-mCherry foci is reported at the indicated time points. Data are from 3 independent biological replicates and shown as the mean ± SD; *n* = 180 cells analyzed in each strain background. Statistical comparisons for the frequencies of cells with 2+ ComEA-mCherry foci were made by one-way ANOVA with Tukey’s multiple comparison test of the log-transformed data. LOD, limit of detection. NS, not significant. ** = *p* < 0.01

Following the uptake of dsDNA into the periplasm, a single strand of this tDNA is translocated across the inner membrane into the cytoplasm via the integral membrane protein ComEC (15, 16). And it is hypothesized that the strand of DNA that is not translocated into the cytoplasm is degraded. The molecular mechanism underlying the translocation of this single-stranded tDNA across the inner membrane remains very poorly understood. However, single cell tracking of the spatiotemporal dynamics of tDNA integration are consistent with the recombination of ssDNA (17). Thus, we hypothesized that SbcB could be facilitating NT by promoting the degradation of one strand of tDNA to help translocate ssDNA into the cytoplasm. So, we next sought to test whether SbcB affects the translocation of tDNA into the cytoplasm.

In *V. cholerae*, the translocation of DNA across the outer and inner membranes is temporally uncoupled (18). Consistent with this, we find that the number of ComEA-mCherry foci in the Δ*comEC* strain following a 30 min incubation with tDNA is indistinguishable from the parent (**Fig. 2B**), indicating that the Δ*comEC* mutant still exhibits normal tDNA uptake into the periplasm. Because uptake across the two membranes is uncoupled, we hypothesized that the translocation of tDNA from the periplasm to the cytoplasm could be tracked indirectly using the same ComEA-mCherry reporter. Because as DNA is translocated across the inner membrane there should no longer be a DNA substrate for ComEA-mCherry to bind to in the periplasm, thus, resulting in the resolution of ComEA-mCherry foci (**Fig. 2A**). For this assay, it was critical to prevent cells from taking up any new tDNA, which would confound the ability to look at the resolution of ComEA-mCherry foci due to DNA translocation across the inner membrane. So after incubating cells with tDNA for 30 mins to allow for tDNA uptake into the periplasm, cells were treated with DNAse to prevent any additional tDNA uptake. Then, cells were incubated for 2 additional hours to allow for DNA translocation across the inner membrane. When we performed this experiment, we saw that the number of cells with ≥2 ComEA-mCherry foci was significantly reduced in the parent but not in the Δ*comEC* mutant. This is consistent with tDNA translocating across the inner membrane in the parent but not in the Δ*comEC* mutant, as expected. In this assay, the Δ*sbcB* mutant was indistinguishable from the parent, suggesting that SbcB does not facilitate the translocation of tDNA across the inner membrane.

### SbcB does not facilitate branch migration during NT

Because SbcB was not required for the translocation of tDNA into the cytoplasm, we hypothesized that it was playing a role in the downstream process of tDNA integration. Our prior work demonstrates that following RecA-dependent strand invasion, the NT-specific helicase ComM facilitates branch migration of tDNA into the genome (19). An unusual property of ComM is that it exhibits bidirectional helicase activity and can translocate either 5’→3’ or 3’→5’ on DNA substrates *in vitro* (19). This bidirectional helicase activity, if stochastic, would be equally likely to promote or inhibit tDNA integration during NT. From experiments tracking the spatiotemporal dynamics of tDNA integration in single cells, however, it is clear that ComM promotes highly efficient homologous recombination during NT (17). This suggests that there may be additional factors *in vivo* that promote ComM directionality. Thus, we hypothesized that the exonuclease activity of SbcB may facilitate NT by regulating the directionality of ComM-dependent branch migration. Specifically, that SbcB degrades the genomic strand displaced by the integrating tDNA so that ComM branch migration can only proceed in one direction.

To test this, we used a previously established assay to assess branch migration activity *in vivo* (19). This assay detects branch migration indirectly via assessing the comigration of two linked genetic markers. Specifically, between a Trimethoprim resistance marker (Tm^R^) and a nonsense mutation (*i*.*e*., a premature stop codon) in *lacZ*, which we refer to as *lacZ*\* (**Fig. 3A**). We used two different tDNA products: one where Tm^R^ and *lacZ*\* are separated by 243 bp, and another where they are separated by 820 bp.

**Fig. 3.**
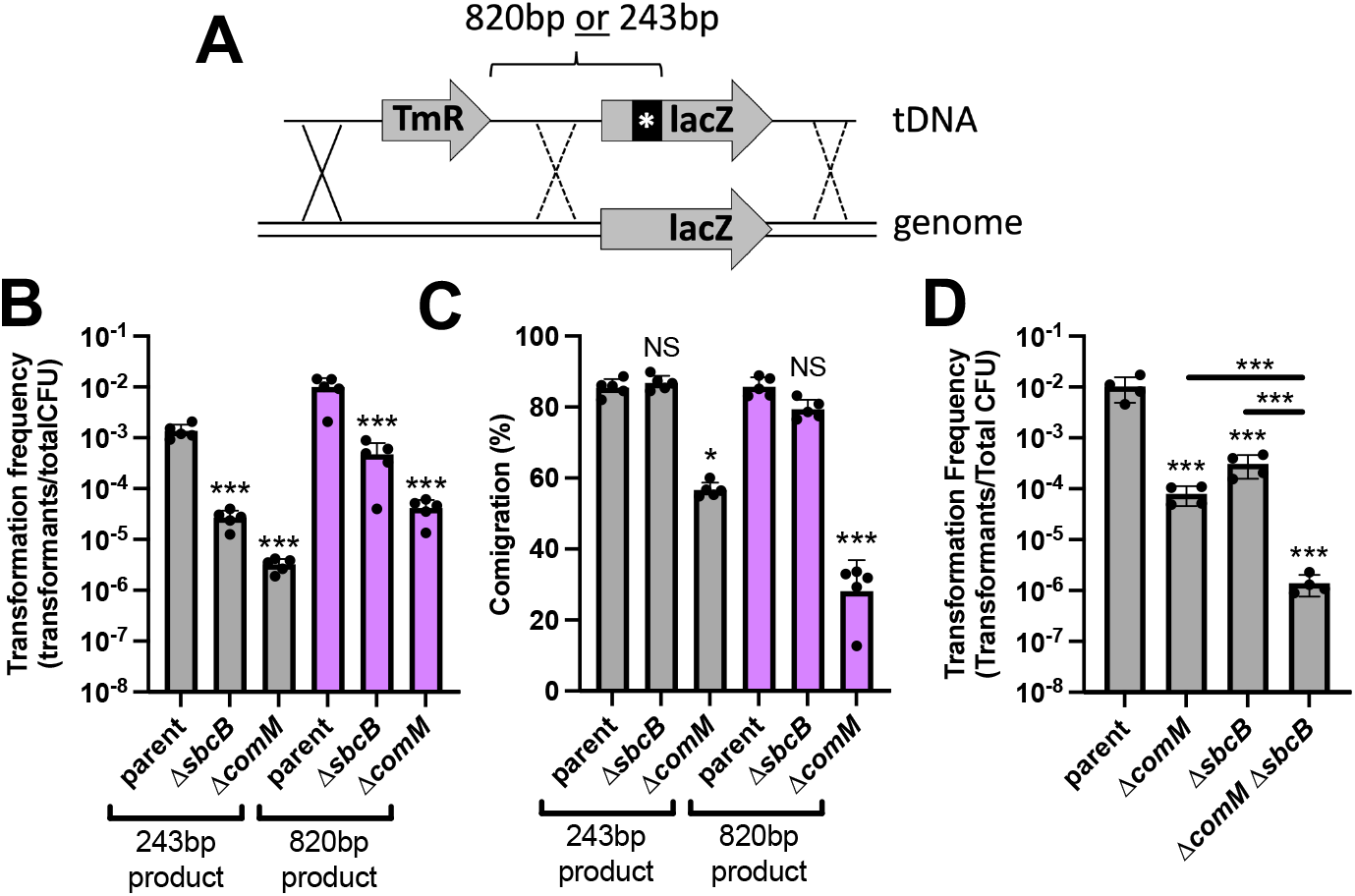
SbcB does not promote ComM-dependent branch migration. (**A**) Schematic of the experimental setup to assess branch migration via linkage analysis between a trimethoprim resistance marker (Tm^R^) and a nonsense mutation in *lacZ*. Two different tDNAs were used: one where the *lacZ* nonsense mutation was 243 bp away from the Tm^R^ cassette, and another where the spacing was 820 bp. The two tDNA products described in **A** were naturally transformed into the indicated strains and the (**B**) transformation frequency and (**C**) comigration frequency (i.e., the frequency of linkage between the Tm^R^ and *lacZ* nonsense mutation) are reported. All strains in these assays contained a Δ*mutS* mutation to prevent mismatch repair from confounding the results. (**D**) Natural transformation assays of the indicated strains. All data are from at least 4 independent biological replicates and shown as the mean ± SD. Statistical comparisons were made by one-way ANOVA with Tukey’s multiple comparison test of the log-transformed data. NS, not significant. *** = *p* < 0.001, ** = *p* < 0.01, * = *p* < 0.05. Statistical identifiers directly above bars represent comparisons to the parent.

Cells are transformed with this linear tDNA and if branch migration is efficient, both markers are integrated into the genome. But if branch migration is inefficient, then cells may integrate the Tm^R^ marker (which is selected for in the experiment), but not the genetically linked *lacZ*\* mutation. When we assess the transformation frequency of the Tm^R^ marker in these experiments, we find that Δ*sbcB* and Δ*comM* are significantly attenuated compared to the parent, which is consistent with SbcB and ComM both playing an important role during NT (**Fig. 3B**). When we assess the associated comigration (*i*.*e*., the linkage with the *lacZ*\* mutation), we find that Δ*comM* is significantly reduced compared to the parent, which is consistent with the established role for ComM in branch migration during NT (**Fig. 3C**). However, the comigration of Δ*sbcB* is indistinguishable from the parent, despite the reduced transformation frequency in this background (**Fig. 3B-C**). This suggests that SbcB may not facilitate branch migration as hypothesized.

To further test this, we assessed the transformation frequency of a Δ*sbcB* Δ*comM* double mutant. If these proteins work together to promote branch migration, we would expect the transformation frequency of this double mutant to be similar to the single Δ*comM* or Δ*sbcB* mutant (*i*.*e*., an epistasic interaction). But if these genes do not act together, then we would expect the transformation phenotype of Δ*sbcB* Δ*comM* to be the sum of the transformation deficits observed in each single mutant. When we performed this experiment using linear tDNA, we find that the reduction in NT in Δ*comM* Δ*sbcB* double mutant is the sum of the transformation defects observed in Δ*comM* and Δ*sbcB* single mutants (**Fig. 3D**). These results further suggest that SbcB and ComM act at distinct steps during NT. Thus, SbcB does not facilitate branch migration during NT.

### SbcB acts upstream of ComM-dependent branch migration

Our data thus far show that SbcB acts downstream of DNA uptake into the cytoplasm. And that SbcB does not work with ComM to facilitate branch migration. To continue to narrow down where SbcB acts during NT, we next sought to determine whether SbcB acts upstream or downstream of ComM-dependent branch migration. We have previously demonstrated that a fluorescent translational fusion of ComM (GFP-ComM) dynamically forms foci in a manner that is dependent on RecA-mediated strand invasion (17). Thus, tracking the formation of GFP-ComM foci demarcates cells that are actively undergoing branch migration during NT (**Fig. 4A**). To test whether SbcB acted upstream of ComM-dependent branch migration, we assessed GFP-ComM focus formation in the parent and Δ*sbcB* by epifluorescence timelapse microscopy. We found that the percentage of cells that exhibited a GFP-ComM focus, and the number of GFP-ComM foci generated in those cells was significantly lower in the Δ*sbcB* mutant compared to the parent (**Fig. 4B-C**). These results suggest that SbcB is acting upstream of ComM-dependent branch migration.

**Fig. 4.**
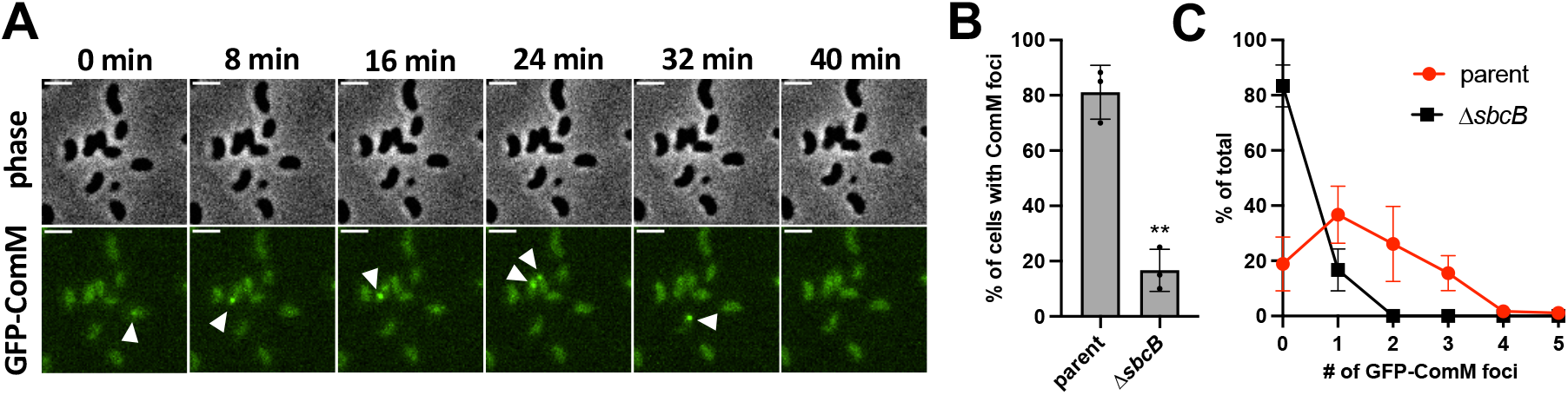
SbcB acts upstream of ComM-dependent branch migration. (**A**) Montage of timelapse imaging of cells expressing a GFP-ComM fluorescent fusion incubated with tDNA. White arrows indicate the appearance of transient GFP-ComM foci. Scale bar, 2 µm. Experiments were performed as in **A** in the parent and Δ*sbcB* background to quantify the (**B**) proportion of cells that exhibited any GFP-ComM foci and the (**C**) distribution of the GFP-ComM foci observed. Data are from 3 independent biological replicates and shown as the mean ± SD; *n* = 180 cells analyzed in each strain background. Statistical comparison in **B** was made by unpaired Student’s t-test of the log-transformed data. ** = *p* < 0.01. Statistical identifier directly above bar represents comparisons to the parent.

### SbcB is epistatic with RecA

Our data suggest that SbcB is acting downstream of DNA uptake into the cytoplasm and upstream of ComM-dependent branch migration. A prerequisite for ComM-dependent branch migration is RecA-mediated strand invasion (17). So next, we sought to determine whether SbcB facilitates RecA-mediated strand invasion. RecA is indispensable for homologous recombination. Thus, Δ*recA* is incapable of NT with linear tDNA that must integrate into the genome (13, 17).

Cells can also take up replicating plasmids by NT, and these products do not require homologous recombination into the host genome (19). As a result, recombination is dispensable when a replicating plasmid is used as tDNA. Consistent with this, Δ*comM* exhibits no reduction in NT with a replicating plasmid as tDNA compared to the parent (**Fig. 5**), which is in stark contrast to when linear tDNA that must recombine into the genome is used (**Fig. 3D**). Also, Δ*recA* is capable of NT with a replicating plasmid as tDNA, although this mutant does exhibit an ∼10-fold reduction in NT (**Fig. 5**). DprA is an essential NT protein that protects ssDNA following translocation into the cytoplasm. Because DprA is critical for NT when a replicating plasmid is used as tDNA, this suggests that cells are still taking up DNA via the same NT pathway in these experiments (**Fig. 5**).

**Fig. 5.**
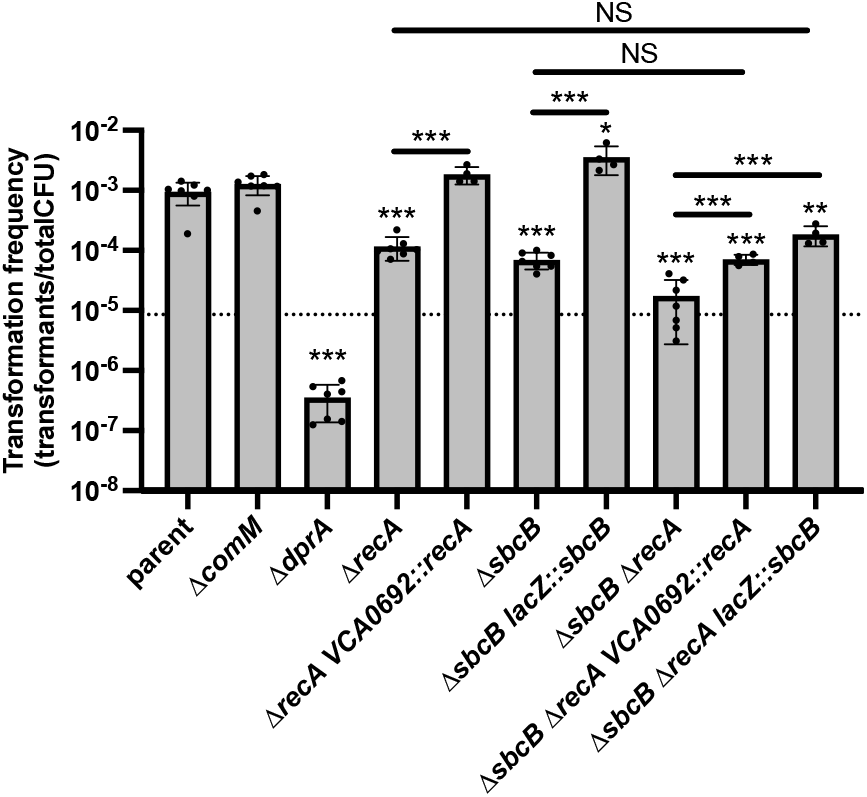
SbcB and RecA facilitate NT when a replicating plasmid is used as the tDNA but are non-epistatic. Natural transformation assays of the indicated strains using a replicating plasmid as the tDNA. Data are from at least 4 independent biological replicate and shown as the mean ± SD. Statistical comparisons were made by one-way ANOVA with Tukey’s multiple comparison test of the log-transformed data. NS, not significant. *** = *p* < 0.001, ** = *p* < 0.01, * = *p* < 0.05. The dotted line represents the expected phenotype for Δ*sbcB* Δ*recA* if these mutations are non-epistatic. Statistical identifiers directly above bars represent comparisons to the parent.

Similar to Δ*recA*, we observed an ∼10-fold reduction in NT in the Δ*sbcB* mutant. Importantly, the phenotypes of Δ*sbcB* and Δ*recA* could be complemented in these assays (**Fig. 5**). To determine if SbcB and RecA work together to promote NT, we assessed the phenotype of the Δ*sbcB* Δ*recA* double mutant. If these proteins do not work together to facilitate NT, we would expect that these mutations are non-epistatic, which would be observed as an NT defect in the double mutant that matches the product of the defects observed in each single mutant. In this case, each mutant exhibits an ∼1-log defect in NT, thus, for a non-epistatic interaction, the double mutant should exhibit an ∼2-log reduction in NT compared to the parent (**Fig. 5**, dotted line). If these proteins do work together, we would expect antagonistic epistasis, which would be observed as an NT defect of <2-log in the double mutant. If these proteins perform the same function (*i*.*e*., act redundantly), we would expect synergistic epistasis, which would be observed as an NT defect of >2-log in the double mutant. When we performed this experiment, we found that the double mutant exhibited an ∼2-log defect in NT, which is consistent with these mutations being non-epistatic (**Fig. 5**). And ectopic complementation of Δ*sbcB* Δ*recA* with either *sbcB* or *recA* restored NT to the level of the corresponding single mutant (**Fig. 5**). Thus, these proteins likely facilitate recombination during NT in distinct ways. While it is not clear why RecA and SbcB are required for NT with a replicating plasmid tDNA (see **Discussion**), the observed reduction in NT in Δ*sbcB* further suggests that SbcB is acting upstream of branch migration to play a presynaptic role during this process.

### SbcB does not require its exonuclease activity to promote NT

SbcB is a DnaQ family exonuclease that has a highly conserved catalytic domain (20). To test whether the exonuclease activity of SbcB was required for its stimulatory effect on NT, we complemented Δ*sbcB* with an ectopic copy of *sbcB* containing mutations to two invariantly conserved residues within the catalytic site (*sbcB*\* = *sbcB*^*D13AE15A*^). When we assessed NT, we were very surprised to find that *sbcB*\* did not phenocopy Δ*sbcB* (**Fig. 6A**). In fact, the NT frequency for Δ*sbcB lacZ*::*sbcB*\* was indistinguishable from the parent (**Fig. 6A**).

**Fig. 6.**
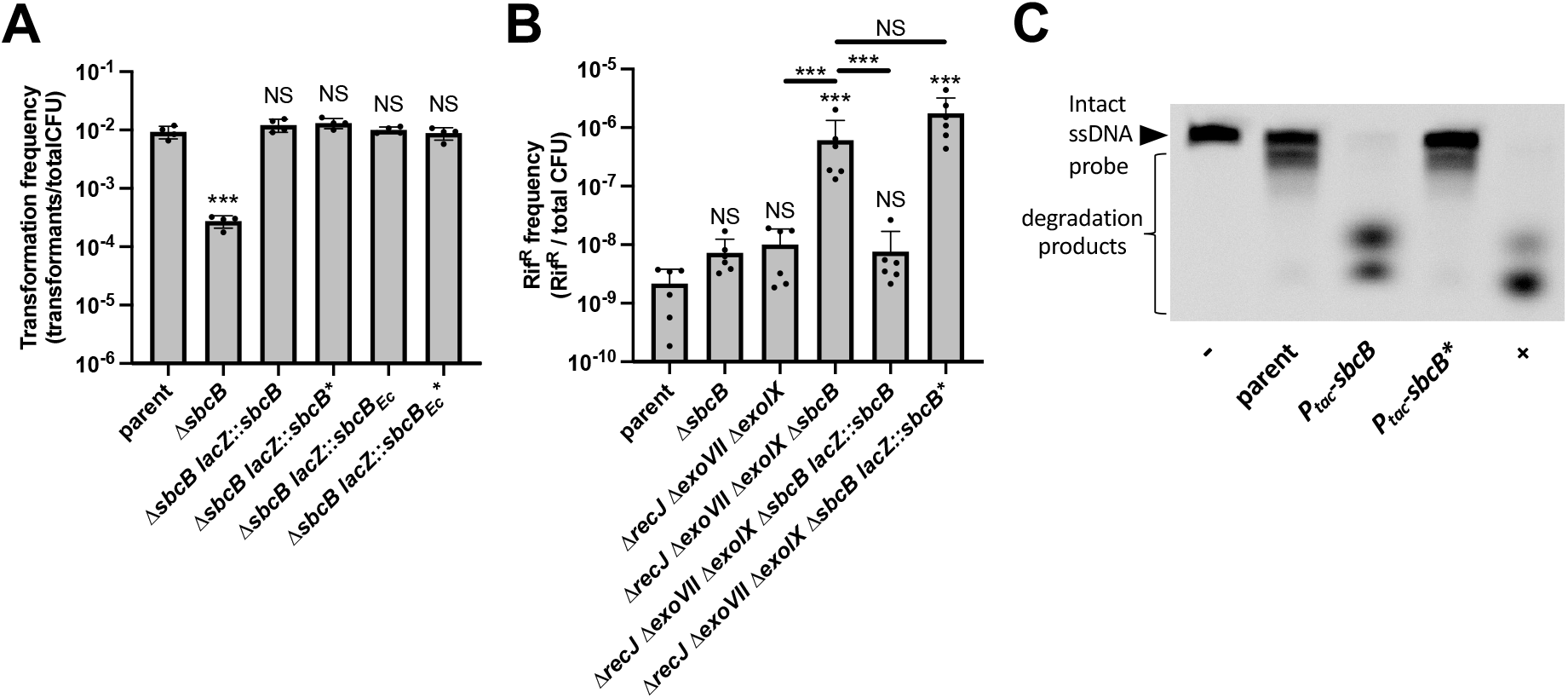
SbcB facilitates NT in *Vibrio cholerae* independent of its exonuclease activity. (**A**) Natural transformation assays of the indicated strains using linear tDNA. (**B**) Quantifying the frequency of spontaneous Rif^R^ mutants in the indicated strain backgrounds. All data in **A** and **B** are from at least 4 independent biological replicates and shown as the mean ± SD. NS, not significant. *** = *p* < 0.001. Statistical identifiers directly above bars represent comparisons to the parent. (**C**) Exonuclease activity assay using cell lysates from the indicated strains. All strains were grown overnight in 1 mM IPTG for these assays. “-” denotes a reaction where no lysate was added, while “+” denotes a sample where purified SbcB (*i*.*e*., NEB Exonuclease I) was added. Data are representative of two independent experiments.

The catalytic residues of SbcB mutated to generate SbcB* are invariantly conserved among DnaQ family exonucleases (20), however, we sought to more rigorously test whether these mutations eliminate exonuclease activity. In *E. coli*, SbcB facilitates mismatch repair by participating in the excision reaction via its exonuclease activity. This is supported by the observation that 4 exonucleases, RecJ, ExoVII, ExoX, and SbcB, are redundant for mismatch repair in *E. coli* where the *recJ exoVII exoX sbcB* quadruple mutant has an ∼6-fold elevated rate of acquiring spontaneous Rif^R^ mutations compared to the *recJ exoVII exoX* triple mutant (21). *V. cholerae* does not have a homolog of ExoX, but it does encode homologs to the ssDNA exonucleases RecJ, ExoVII and ExoIX. So first, we assessed whether SbcB facilitates mismatch repair in *V. cholerae*. To do this, we determined the spontaneous Rif^R^ mutation rate in a Δ*recJ ΔexoVII ΔexoIX* triple mutant vs a Δ*recJ ΔexoVII ΔexoIX ΔsbcB* quadruple mutant. We found that the latter strain exhibited a spontaneous Rif^R^ rate that was ∼50-fold higher (**Fig. 6B**). However, the spontaneous Rif^R^ rate for the Δ*sbcB* single mutant was indistinguishable from the parent (**Fig. 6B**). This suggests that SbcB is redundant with other exonucleases for mismatch repair in *V. cholerae*, as previously observed in *E. coli* (21). This redundancy with other exonucleases also suggests that it is likely the exonuclease activity of SbcB that is required for mismatch repair. When we ectopically expressed *sbcB* vs *sbcB*\* in the Δ*recJ ΔexoVII ΔexoIX* Δ*sbcB* mutant, we found that the strain complemented with wildtype *sbcB* reduced the mutation rate to the level of *recJ exoVII exoX*, while the strain with *sbcB*\* was indistinguishable from the Δ*recJ ΔexoVII ΔexoIX ΔsbcB* parent (**Fig. 6B**). This strongly suggests that the mutations in SbcB* eliminate its exonuclease activity as hypothesized.

We also assessed ssDNA exonuclease activity more directly to further test whether the mutations in SbcB* eliminate its exonuclease activity. For these assays, we generated P_*tac*_-overexpression constructs for SbcB and SbcB*. Then, we incubated cell lysates from these strains with a 5’-end labeled ssDNA oligonucleotide. In the presence of P_*tac*_-*sbcB* lysate, the ssDNA probe is rapidly degraded when compared to lysate from a parent strain lacking an SbcB overexpression construct (**Fig. 6C**). And the degradation products formed by the P_*tac*_-*sbcB* lysate resemble the degradation products formed by commercially available purified *E. coli* SbcB (**Fig. 6C**). Degradation of ssDNA by P_*tac*_-*sbcB*\* lysates, however, are indistinguishable from the parent lysate. All together, these data demonstrate that the mutations used to generate SbcB* eliminate its exonuclease activity.

SbcB is narrowly conserved among gammaproteobacteria (22), and it has predominantly been studied and characterized in the model organism *E. coli*. SbcB from *V. cholerae* is well-conserved to its *E. coli* homolog (59% identity, 74% similarity). To test whether *V. cholerae* SbcB has unique properties that allow it to facilitate NT, we complemented the Δ*sbcB* mutant by ectopically expressing *E. coli sbcB* (*sbcB*_*Ec*_). We found that SbcB_Ec_ could fully complement the Δ*sbcB* mutant as it restored NT back to parent levels (**Fig. 6A**). Furthermore, a catalytic mutant of SbcB_Ec_ (*sbcB*_*Ec*_* = *sbcB*_*Ec*_ ^*D15AE17A*^) also fully restored the NT of Δ*sbcB* back to parent levels (**Fig. 6A**). All together, these results indicate that the exonuclease activity of SbcB is not required to facilitate NT, and that this is activity is not a unique property of *V. cholerae* SbcB.

### SbcB interacts with RecA

Our data suggest that SbcB is likely facilitating RecA-dependent strand invasion in a manner that does not rely on its exonuclease activity. One possibility is that SbcB directly interacts with RecA to help coordinate its activity and promote recombination. To test this, we performed <u>B</u>acterial <u>A</u>denylate <u>C</u>yclase <u>T</u>wo <u>H</u>ybrid (BACTH) assays (23) to assess the interaction between SbcB* and RecA. When we performed these assays, we found that SbcB* exhibited interactions with itself and RecA (**Fig. 7**). Because SbcB and RecA can both bind to ssDNA, this interaction may not reflect a direct interaction between these proteins, but could instead represent an interaction that is bridged by a ssDNA substrate in the cell. To test this, we also performed BACTH analysis with DprA, which is also a ssDNA-binding protein (5). If ssDNA was helping to bridge interactions in our BACTH assays, we hypothesized that we should also observe interactions between SbcB* and DprA. However, when we tested this, no interactions were observed between SbcB* and DprA (**Fig. 7**). This suggests that the interactions observed between SbcB* and RecA are specific.

**Fig. 7.**
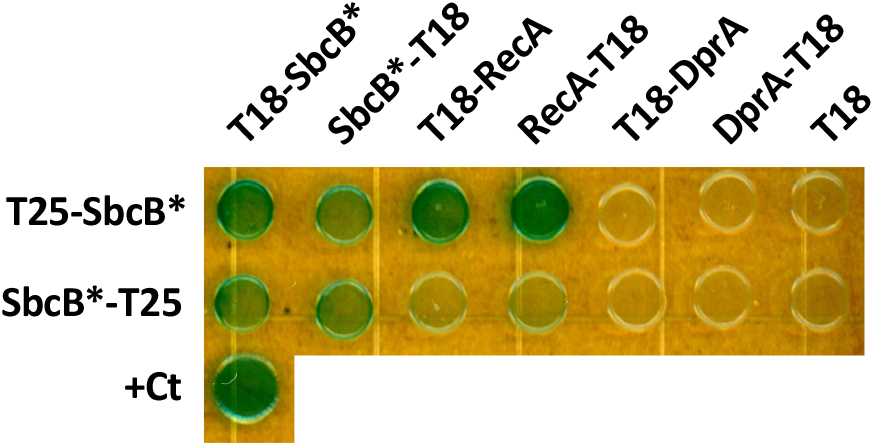
SbcB interacts with RecA. Bacterial Adenylate Cyclase Two-Hybrid (BACTH) assays were performed to assess pairwise interactions between SbcB* (SbcB* = the SbcB^D13A,E15A^ catalytic mutant) and RecA or DprA. The indicated proteins were fused to the T18 or T25 fragments of adenylate cyclase as indicated. Leucine zipper fusions to T18 and T25 served as a positive control for interacting proteins (+Ct). Data are representative of 3 independent biological replicates.

## DISCUSSION

Our results demonstrate that SbcB acts downstream of tDNA uptake and upstream of ComM-dependent branch migration. Also, they show that the exonuclease activity of SbcB is dispensable for its role in promoting NT. These results help to narrow down the potential role that SbcB plays during NT.

Prior work demonstrates that mutant alleles of SbcB with diminished exonuclease activity can protect 3’ ends (24, 25). Also, single-stranded binding protein (SSB) directly interacts with SbcB to promote its exonuclease activity (26). During NT, tDNA is coated with DprA, which can evict SSB and promote the loading of RecA (5). So, SSB may be excluded from interacting with SbcB at the 3’ end of the tDNA, which may hamper SbcB exonuclease activity during NT *in vivo*. Thus, SbcB may help protect the 3’ end of tDNA to facilitate recombination during NT. Our observation that an exonuclease defective allele of SbcB is equally capable of promoting NT is consistent with this hypothesis.

Our data demonstrate that SbcB and RecA both contribute to NT when a replicating plasmid is used as the tDNA. Because ssDNA is translocated into the cytoplasm during NT, cells must take up multiple independent complementary tDNA molecules to reassemble the plasmid into a replicative form. In the simplest case, cells would require the uptake of 2 tDNA molecules: one molecule that represents the entire ssDNA sequence of the plasmid, and a second partially complementary molecule that can prime synthesis of the other strand. While RecA-dependent strand invasion would not be necessary for this assembly, it is possible that RecA promotes annealing of complementary sequences, resulting in the observed ∼10-fold deficit for Δ*recA* when a replicating plasmid is used as tDNA. Our data also showed an ∼10-fold deficit for Δ*sbcB* in these assays. However epistasis analysis indicated that these mutations were non-epistatic. Thus, RecA and SbcB likely facilitate NT with a replicating plasmid in distinct ways. RecA may facilitate annealing, while SbcB helps protect the 3’ end of the tDNA.

RecA assembly on ssDNA occurs unidirectionally from 5’ → 3’ (27). Thus, for RecA to bind to the 3’ end of a long tDNA molecule, it must be processively assembled from an upstream 5’ site. As a 3’ → 5’ exonuclease, SbcB binds to the 3’ ends of ssDNA molecules. Our results provide preliminary genetic evidence that SbcB directly interacts with RecA, which may help recruit RecA to the 3’ end of single-stranded tDNA. When a linear tDNA homologous to the host chromosome is used, RecA loading on the 3’ end could also facilitate homologous recombination by promoting strand invasion of the 3’ end into the genome. This could allow the integrated 3’ end to initiate DNA replication, which would stabilize the recombination intermediate (28). Interestingly, prior data suggest that SbcB facilitates RecA-dependent recombination *in vitro* (29). In these studies, however, it was presumed that this stimulatory effect of SbcB was due to its exonuclease activity.

SbcB has primarily been characterized in *E. coli*, and a number of studies have noted distinct recombination phenotypes in exonuclease defective alleles of *sbcB* (*i*.*e*., *sbcB15*) vs *sbcB* null mutants (*i*.*e*., Δ*sbcB* and *xonA2*) (24, 30, 31). These studies suggest that SbcB has an activity that facilitates recombination beyond its exonuclease activity. Consistent with what is observed in *E. coli*, we uncover a phenotype for SbcB in *V. cholerae* that is distinct from its exonuclease activity. Interestingly, revealing these phenotypes for SbcB in *E. coli* relied on genetic backgrounds that lack other recombination proteins like RecBCD and SbcCD (24, 30, 31), while in *V. cholerae* we observe this phenotype even when these recombination factors are intact. Regardless, these results suggest that an exonuclease independent activity of SbcB may be conserved. Indeed, this is supported by our observation that *E. coli* SbcB can complement the NT defect of a *V. cholerae* Δ*sbcB* mutant. It is tempting to speculate that this exonuclease-independent activity is due to SbcB-RecA interactions, which we provide preliminary genetic evidence for here.

All together, these results elucidate a new role for the canonical exonuclease SbcB in facilitating the integration of tDNA during horizontal gene transfer by NT in *V. cholerae*. Future work will focus on testing on testing SbcB-RecA interactions biochemically and the impact of these interactions on recombination both *in vitro* and *in vivo*. While NT is broadly conserved in diverse microbial species (32), SbcB is narrowly conserved among gammaproteobacteria (22). So it is tempting to speculate that the role that SbcB plays during NT in *V. cholerae*, is facilitated by other recombination proteins in more distantly related naturally transformable species.

## METHODS

### Bacterial Strains and Culture Conditions

All strains were derived from *V. cholerae* E7946 (33). Cells were routinely grown in LB Miller broth and agar supplemented with spectinomycin (200 µg/mL), kanamycin (50 µg/mL), trimethoprim (10 µg/mL), tetracycline (0.5 µg/mL), sulfamethoxazole (100 µg/mL), chloramphenicol (1 µg/mL), erythromycin (10 µg/mL), carbenicillin (20 µg/mL), or zeocin (100 µg/mL) as appropriate.

### Construction of mutant strains

All strains were generated by natural transformation / MuGENT using splicing-by-overlap-extension (SOE) PCR mutant constructs exactly as previously described (34, 35). Ectopic expression constructs were generated by SOE PCR and integrated onto the *V. cholerae* genome exactly as previously described (36). All strains were confirmed by PCR and/or sequencing. For a detailed list of all mutants used in this study, see **Table S1**. For a complete list of primers used to generate mutant constructs, see **Table S2**.

### Natural transformation assays

As discussed in the text, cells were engineered to bypass the native signals required for NT in *V. cholerae*. Specifically, the master competence regulator TfoX was overexpressed via an isopropyl β-d-1-thiogalactopyranoside (IPTG)-inducible P_*tac*_ promoter, and the cells are genetically locked in a high cell density state via deletion of *luxO*, as previously described. Chitin-independent transformation assays to determine NT frequency were performed exactly as previously described (1). Briefly, cells were grow at 30ºC rolling in LB supplemented with 100 µM IPTG, 20 mM MgCl_2_, 10 mM CaCl_2_ to late-log. Then, ∼10^8^ CFUs were diluted into instant ocean medium (7g/L; Aquarium Systems). Next, transforming DNA (tDNA) was added into each reaction as appropriate. For linear tDNA that must integrate into the genome, 100 ng of a ΔVC1807::Erm^R^ product was used. VC1807 is a frame-shifted transposase gene. For replicating plasmid as tDNA, pBAD18-Kan (37) was miniprepped from *E. coli* TG1. Control reactions were performed for each strain where no tDNA was added. Cells were incubated with tDNA overnight. Then, reactions were outgrown by adding 1mL of LB and shaking at 37ºC for 3 hours. Reactions were then plated for quantitative culture by dribble plating on LB agar supplemented with the appropriate antibiotic to quantify transformants, and plain LB to quantify total viable counts. Transformation frequency is defined as the CFU/mL of transformants divided by the CFU/mL of total viable counts.

For assays where branch migration was indirectly assessed via linkage analysis, transformation reactions were performed exactly as described above except that they were plated onto LB agar supplemented with trimethoprim and X-gal (40 µg/mL). On these plates, all cells integrated the Tm^R^ marker, and if cells integrated the genetically linked *lacZ* nonsense mutation they would make white colonies. If they did not integrate the linked lacZ nonsense mutation, they would make blue colonies on these plates. Thus, the comigration frequency is defined as the percent of Tm^R^ colonies that were white on these plates.

### Spontaneous Rif^R^ mutant frequency assays

All strains were isolation streak plated onto plain LB agar. Then, single colonies were picked and grown overnight in plain LB. These cultures were then concentrated and ∼10^10^ CFUs were spread onto LB agar plates supplemented with Rifampicin 100 µg/mL to quantify Rif^R^ mutants. Concentrated cultures were also dilution dribble plated onto plain LB agar plates to quantify total CFU. The Rif^R^ mutant frequency is defined as Rif^R^ CFU / total CFU.

### Microscopy

Phase contrast and fluorescence images were collected on a Nikon Ti-2 microscope using a Plan Apo × 60 objective, a YFP and/or mCherry filter cube, a Hamamatsu ORCA Flash 4.0 camera and Nikon NIS Elements imaging software.

To assess ComEA-mCherry localization upon DNA uptake, cells were grown to induce competence as described above for natural transformation assays. Then, ∼10^7^ cells were mixed with ∼750 ng of a 6kb PCR product as tDNA. Reactions were incubated at 30ºC static for 30 mins to allow for tDNA uptake into the periplasm. Half the reaction was then placed on ice for imaging at this time point. DNAse was added to the other half of the reaction, which was then incubated at 37ºC for 120 mins. Then, both samples (30 min and 150 min) were placed under 0.2% gelzan pads and imaged. The number of ComEA-mCherry foci was determined by analyzing 60 cells per replicate.

To assess GFP-ComM focus formation, cells were grown to induce competence as described above for natural transformation assays. Then, ∼10^7^ cells were mixed with ∼650 ng of a 6kb tDNA product that will integrate into the genome, and placed under an 0.2% gelzan pad. Samples were then subjected to timelapse imaging where images were taken every 2 min for 3 hours. The number of GFP-ComM foci was determined by analyzing 60 cells per replicate.

### ssDNA exonuclease activity assays

Strains were grown overnight in LB supplemented with 1 mM IPTG. Then, cells were washed and concentrated to an OD_600_ = 25.0 in Buffer A: 10mM Tris pH 7.5, 100 mM NaCl. Then, the following reagents were added to the indicated final concentrations to lyse cells: 1x FastBreak cell lysis reagent (Promega), 100 μg/mL lysozyme, 1 mM PMSF, 1x protease inhibitor cocktail, and 10 mM beta-mercaptoethanol. The 100x inhibitor cocktail contained the following: 0.07 mg/mL phosphoramidon (Santa Cruz), 0.006 mg/mL bestatin (MPbiomedicals/Fisher Scientific), 1.67 mg/mL AEBSF (Gold Bio), 0.07 mg/mL pepstatin A (DOT Scientific), 0.07 mg/mL E64 (Gold Bio) suspended in DMSO). Next, samples were subjected to bead beating with lysing matrix B on a FastPrep-24 instrument (MP Bio) with 3 × 40 s pulses at 6.0 m/s. Lysates were centrifuged at 18,000 x g for 2 mins and the supernatant was transferred to a fresh eppi. The ssDNA probe used was PP327 (kindly provided by Payel Paul and Julia van Kessel), a 5’ IR800CW end-labeled oligonucleotide (see **Table S2**). Next, 20 µL ssDNA exonuclease reactions were setup as follows: 3 µL PP327 (2 µM), 2 µL 10X Exonuclease I reaction buffer (NEB), 13 µL water, and 2 µL of cell lysate. Negative control reactions were performed where lysate was substituted with 2 µL of Buffer A. And a positive control reaction was performed where lysate was substituted with 2 µL (20 U) of purified Exonuclease I (NEB). Reactions were allowed to incubate for 1 min at room temperature. Then, 2 µL of each sample was mixed with 8 µL of 1.25X formamide loading buffer (90% formamide, 37.5 mM EDTA pH 8.0, 2 mg/mL orange G) to stop reactions. Next, samples were electrophoretically separated on a prerun Tris Borate EDTA (TBE)-Urea gel (15% 29:1 acrylamide/bis-acrylamide; 8M Urea) in 1X TBE at 200 V. Gels were then imaged on a Protein Simple FluorChem R instrument on the infrared setting to visualize the IR800 labeled probe.

### Bacterial Adenylate Cyclase Two-Hybrid (BACTH) Assays

Gene inserts were amplified by PCR and cloned into the pUT18C, pUT18, pKT25, or pKNT25 vectors to generate N- and C-terminal fusions to the T18 and T25 fragments of adenylate cyclase. Miniprepped vectors were then cotransformed into *E. coli* BTH101 and outgrown with 500 μL LB for 1 hour at 37°C with shaking. Reactions were plated on LB agar plates supplemented with kanamycin 50 μg/mL and carbenicillin 100 μg/mL to select for transformants that received both plasmids. Single colonies were then picked and grown statically at 30°C in LB supplemented with kanamycin 50 μg/mL and carbenicillin 100 μg/mL. Then, 3 μL of each strain was spotted onto LB agar plates supplemented with kanamycin 50 μg/mL, carbenicillin 100μg/mL, 0.5 mM IPTG and 40 μg/mL X-gal. Plates were incubated at 30°C for 24 hours prior to imaging on an Epson Perfection V600 scanner.

### Statistics

Statistical differences were assessed using GraphPad Prism software. The statical tests employed are indicated in figure legends. Descriptive statistics for all samples, and a comprehensive list of statistical comparisons can be found in **Dataset S1**.

## ACKNOWLEDGEMENTS

This work was supported by grant R35GM128674 from the National Institutes of Health to ABD.

---

**Table S1.**
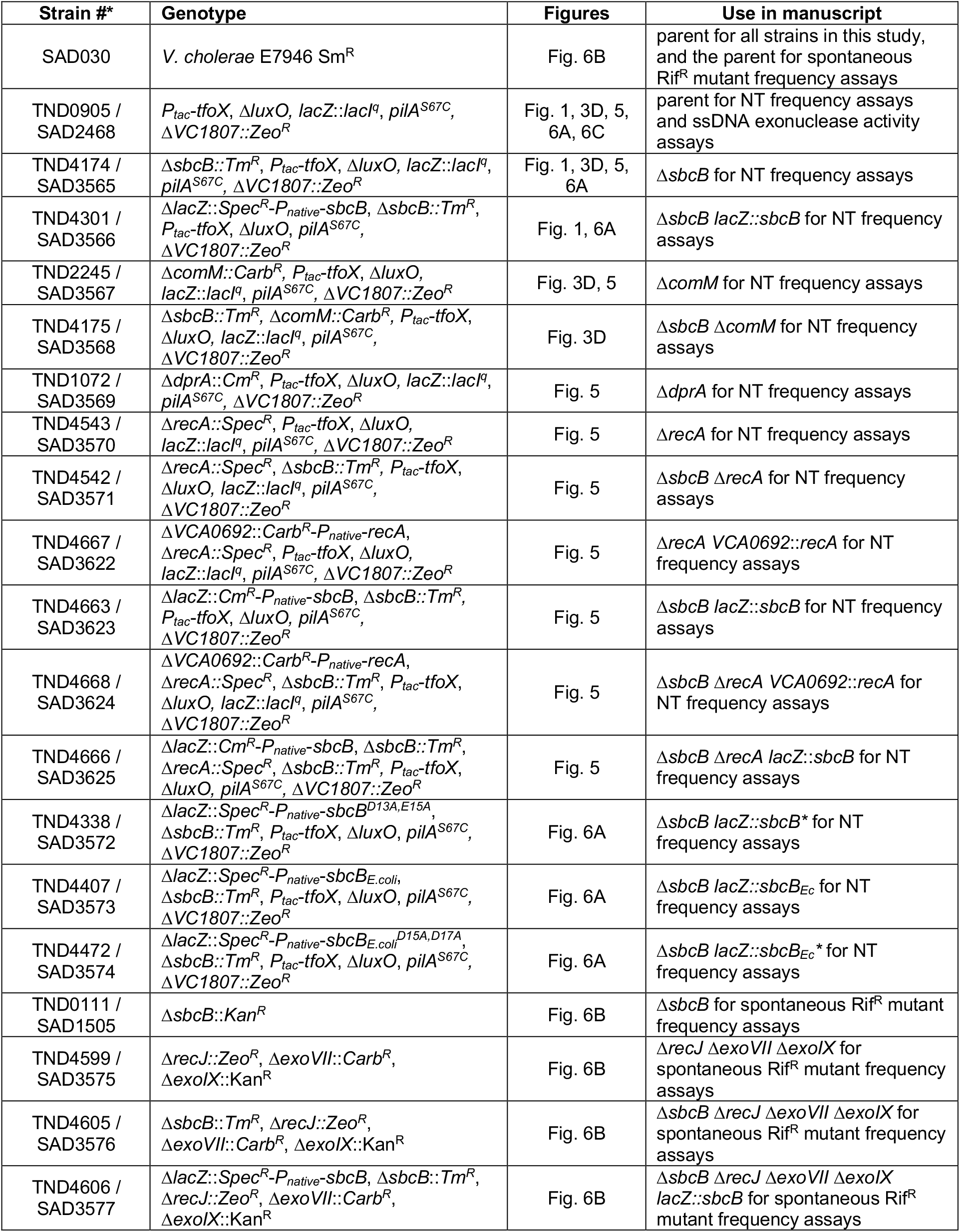
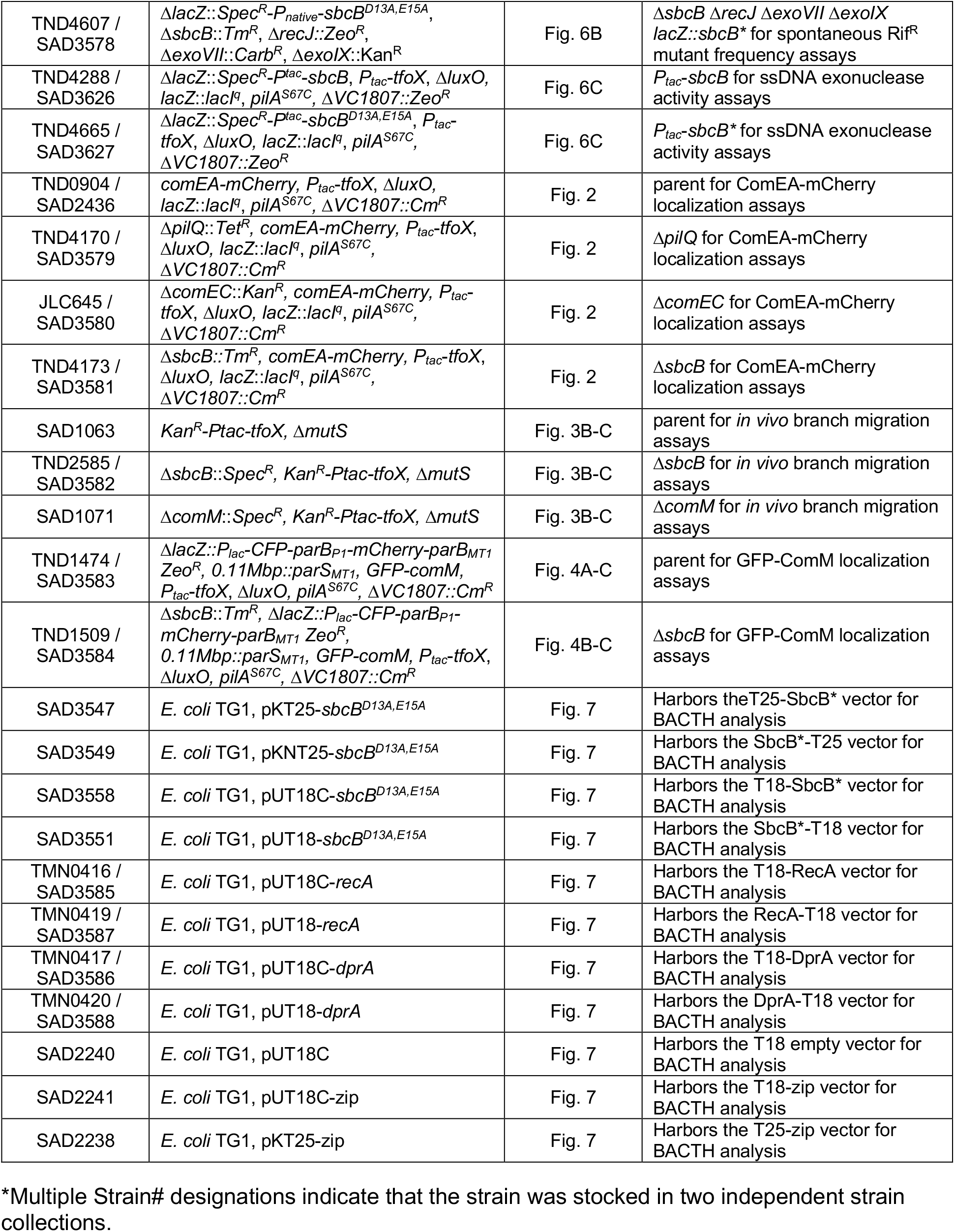
Strains used in this study.

**Table S2.**
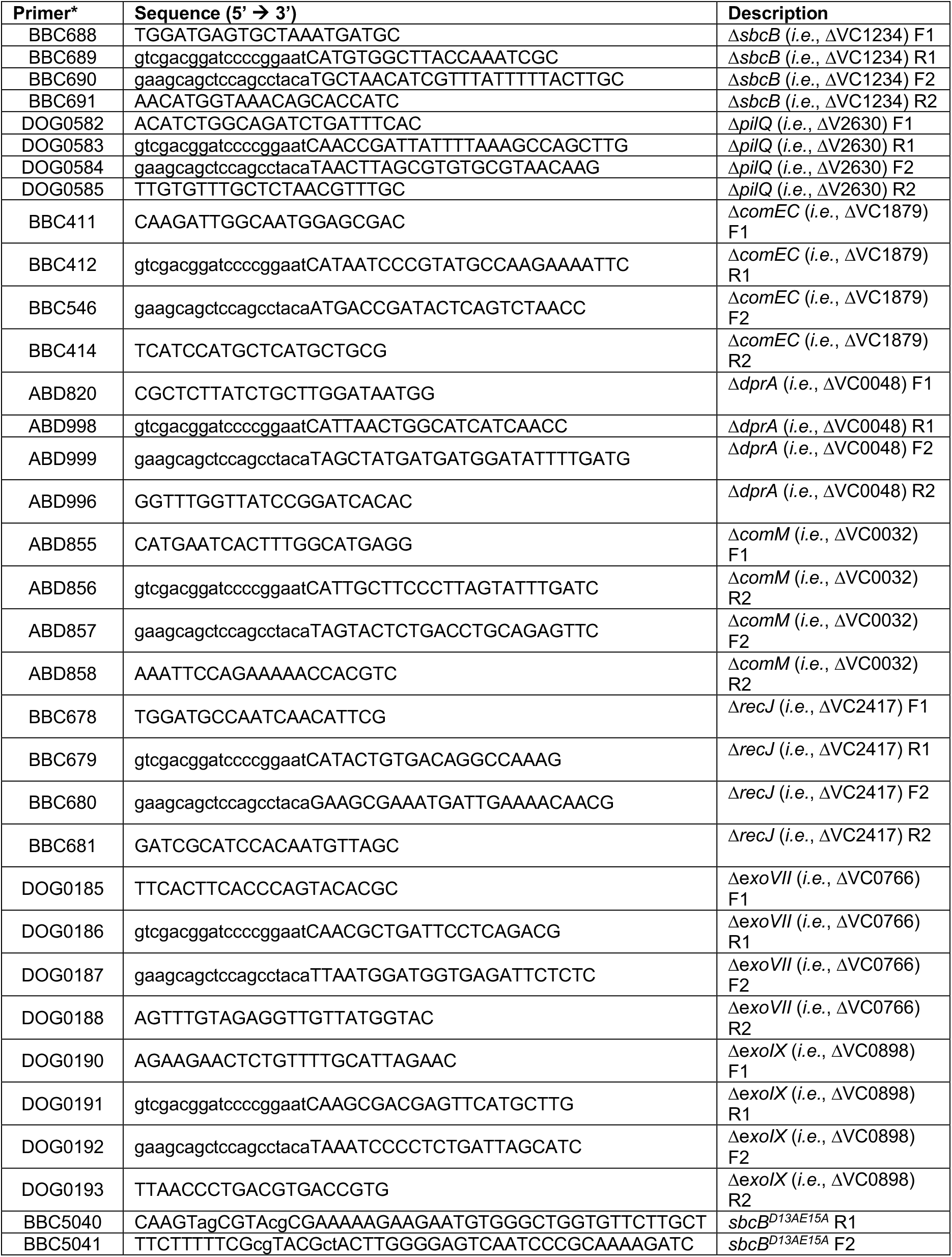
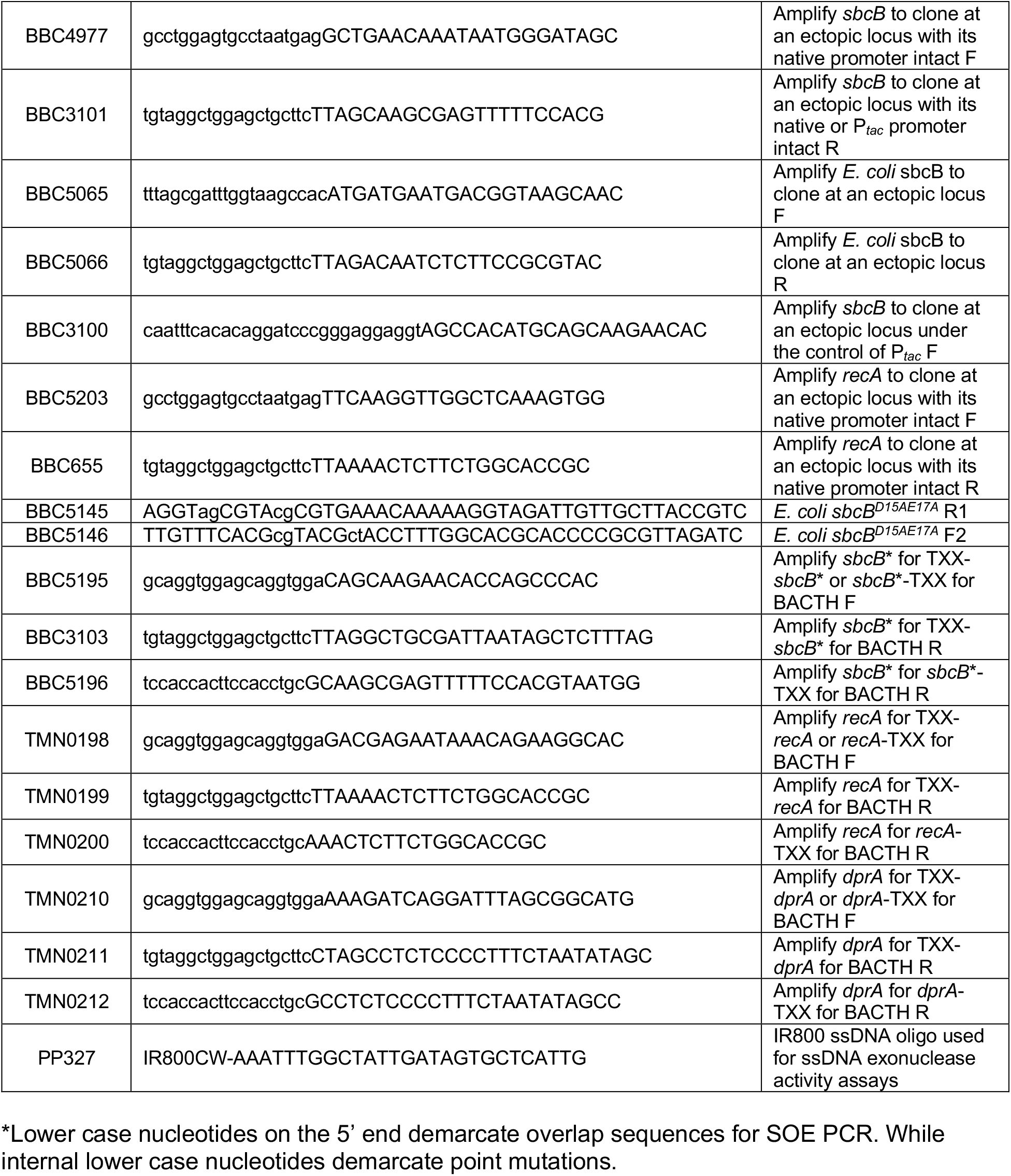
Primers used in this study.

